# Bisphenol A and its Analogues Alter Appetite Control in Zebrafish

**DOI:** 10.1101/2024.04.17.589982

**Authors:** Silvia Karim, Maria Bondesson

**Author notes:** **Corresponding author** Address: Multidisciplinary Engineering and Sciences Hall, 2401 N Milo Sampson Ln, Bloomington, IN 47408-1368.

## Abstract

The regulation of appetite is of growing interest due to the significant rise in global obesity rates. Hunger and satiety are controlled by two hormones with functional activity in the brain; leptin, which is produced in adipocytes and suppresses food intake, and ghrelin, which is produced and released mainly by the stomach and functions as an appetite-stimulatory signal. In this study, zebrafish-based *in vivo* assays were used to examine whether BPA and five of its analogues, BPAF, BPE, BPC, BPC-CL, and BPS affect appetite regulation. The effect of bisphenol exposure on eating behavior was first examined. Four to six days old zebrafish larvae were exposed to a concentration range of the bisphenols and 17β-estradiol, followed by being fed a stained egg yolk powder at day six. After an hour of feeding, the feed in the gut was imaged by microscopy. Quantitative PCR was used to analyze the gene expression of *leptin* and *ghrelin*, as well as eleven other genes involved in appetite control. Exposures to BPA, BPAF, BPE, BPC, BPC-Cl and BPS, resulted in increased amounts of feed in the gut of the larvae in a concentration dependent manner. The qPCR results suggested that *leptin* mRNA expression was downregulated with the increasing concentrations of BPA, BPAF and BPC-Cl, whereas *ghrelin* mRNA expression was upregulated. The expression of several additional anorexigenic genes were downregulated by BPAF and BPC-Cl exposure, whereas orexigenic genes were upregulated. In conclusion, bisphenol exposures resulted in an increased eating behavior in zebrafish larvae, which correlated to increased mRNA expression of appetite-stimulatory genes and decreased expression of satiety-inducing genes. In addition, the results suggest that zebrafish larvae can be used to efficiently assess obesogenic capacity of environmental pollutants.

## Introduction

Both obesity and the related metabolic syndrome are on the rise in many parts of the world. Currently, 17% of American children and adolescents are obese, with a body mass index (BMI) at or above 30 (1). It is anticipated that 86% of Americans will be overweight, with a BMI at or over 25, by 2030. A higher risk of adverse medical consequences is linked to obesity such as heart disease, high blood pressure, diabetes, obstructive sleep apnea, high blood cholesterol and certain types of cancers (2). In the United States, $147 billion or 9% of total medical costs were spent on treating disorders linked to obesity in 2008 (3). Preventing obesity is crucial since obesity is difficult to treat as several physiological and psychological factors hinder weight loss (4, 5)

Obesity is caused by a long-term positive energy balance, thus by more food intake and less energy expenditure. In both humans and rodents, the hypothalamus is a major player in controlling hunger and food intake through interactions between the hormones ghrelin and leptin (6). The “satiety hormone” leptin is secreted by adipose cells. One of its functions is to reduce appetite by stimulating the pro-opiomelanocortin (POMC) neurons in the arcuate nucleus (ARC) (7), which in turn inhibit the action of Agouti-related protein (AgRP) neurons via an opioidergic mechanism (8). Activation of POMC neurons also trigger alpha-melanocyte stimulating hormone (α-MSH) production and release from POMC axon terminals, which in turn activates the anorexigenic melanocortin receptor 4 (MC4R) in the paraventricular nucleus (PVN) (9). The mesoaccumbal dopamine neurons, which are responsible for motivating behaviors such as eating, are also inhibited by leptin (10, 11).

Intestinal enteroendocrine cells generate glucagon-like peptide-1 (GLP-1) upon eating. Similar to leptin, GLP-1 inhibits food intake by interacting with a network of targets, such as Nucleus Tractus Solitarius (NTS) and ARC (12). Another anorexigenic hormone that reduces appetite is peptide YY (PYY), which also is produced by enteroendocrine cells. Recent research has revealed the direct PYY target to be the dorsal raphe nucleus (13).

The “hunger hormone,” ghrelin, is released by the stomach, with levels rising before a meal and decreasing after eating. Ghrelin drives AGRP neuron activity by activating glutamatergic afferents on AGRP neurons (8, 14-16). AGRP neurons, in turn, antagonize the effect of alpha-MSH on MC4R (17). Exogenous administration of ghrelin or the orixogenic neuropeptide Y (NPY) to rats results in stimulation of feeding (18). Release of adrenocorticotrophic hormone (ACTH) is stimulated after ghrelin administration.

Endocrine disrupting chemicals (EDCs) are environmental compounds that interfere with the functions of hormones. A subclass of endocrine disruptors, so called obesogens, are chemicals that alter regulatory processes in metabolism and adipocyte function, resulting in imbalances in the regulation of body weight (19). Obesogens act by altering adipocyte counts, lipid storage, lipid balance, or the control of hunger and satiety. One such chemical is BPA. Human exposure to BPA has been linked to adverse health effects including obesity (20). In rodents, fetal BPA exposure causes the size of adipocytes increase with age, resulting in a larger body fat percentage (21). Also,*in vitro* cell culture data suggest that BPA contributes to the pathogenesis of obesity by promoting adipogenesis, lipid and glucose dysregulation, and inflammation of adipose tissue (22-24). We and others have shown that BPA is a potent estrogen receptor ligand *in vitro* in cell lines and *in vivo* in zebrafish (25-28). BPA also represses thyroid hormone and androgen pathways ((29), reviewed in (30)).

Because BPA is a potent endocrine disruptor, BPA analogues have been synthesized as replacement molecules for BPA. According to a report from the Environmental Protection Agency’s (EPA’s) and National Toxicology Program’s (NTP’s) high-throughput Tox21/ToxCast screening programs, 24 BPA analogues have been synthesized as substitute molecules for BPA (31). Five of these BPA analogues, bisphenol AF (BPAF), bisphenol E (BPE), bisphenol C (BPC), chlorinated bisphenol C (BPC-Cl) and bisphenol S (BPS) are the focus of this study.

Zebrafish (*Danio rerio*) is an efficient *in vivo* model to study obesogenic activity of compounds (32). It is a powerful model to visualize biological effects and chemical metabolism and simplify high-throughput screening. Here, we have utilized the zebrafish model to investigate the obesogenic potential of BPA and its analogues. In an eating assay, we measured the amount of feed that larvae ate after bisphenol exposure. We further performed qPCR to analyze the expression of the hunger and satiety hormones *leptin* and *ghrelin* and their downstream effectors *in vivo* in larvae and juvenile fish.

## Material and Methods

### Chemicals

The chemicals 17β-estradiol (E2) (CAS no. 50-28-2), bisphenol A (CAS no. 80-05-7), bisphenol AF (CAS no. 1478-61-1), bisphenol E (CAS no. 2081-08-5), bisphenol C (CAS no. 79-97-0), bisphenol C-Cl (CAS no. 14868-03-2) and bisphenol S (CAS no. 80-09-1) were bought from Sigma-Aldrich. Compounds were dissolved in dimethyl sulfoxide (DMSO, Fisher Chemical) as 10 mM stock solutions. Paraformaldehyde was bought from Invitrogen.

### Zebrafish treatment for Oil Red O staining

Zebrafish were reared and maintained according to the protocols approved by the Institutional Animal Care and Use Committee at Indiana University Bloomington. The adult fish were housed in automated tank systems supplied with circulating filtered water at 27-28°C on 14h/10h light/dark cycles. The adult fish were fed with Tetramin flakes in the morning and brine shrimp (Artemia, Shrimp Direct) in the afternoon. The fish line used for this study was Gateways (GW) 57A (33) crossed to mitfa^*b692/b692*^ (Nacre; Zebrafish International Resource Center (ZIRC)) to facilitate visualization of feed in the gut without obstruction from melanophores, and without having to treat with 1-phenyl 2-thiourea (PTU).

Adult zebrafish were set up for spawning in the afternoon and embryos collected the morning after and cultured in a Petri dish in E3 embryo medium (0.17 mM KCl, 0.33 mM CaCl2, 0.33 mM MgSO4, 5 mM NaCl, pH 7.0) at 28.5 °C with 14:10 light/dark cycles.

Samples of 12 zebrafish larvae per well were set up in 12-well plates (cellstar, Greiner Bio-One) in 1.2 mL E3. At 4 days post fertilization (dpf) and again at 5 dpf, 17β-estradiol, BPA and the BPA analogues were added in a concentration range. Vehicle only was used as a negative control and did not exceed 1% DMSO.

On day 6, the larvae were fed 2 μL an Oil-Red-O (ORO, Sigma-Aldrich) stained egg yolk powder mixture for 1 h. The egg yolk mixture contained 100 mg egg yolk powder in 0.5 mL ORO and 1 mL E3. After an hour of feeding, the samples were fixed with 4% paraformaldehyde (Invitrogen) overnight. The next day, the samples were imaged under 5X objective using Leica DMi8 (Leica Microsystems) fluorescence microscope equipped with Leica LASX software and the Leica DFC9000 GT camera. The ORO stain in the gut was captured by fluorescence in the red (RFP) filter. At least three biological replicates were performed per compound and concentration.

### Quantification of fluorescence in images and statistical analysis

The fluorescence in images of feed intake in the gut were quantified by the Fiji/imageJ (https://imagej.net/Fiji and https://imagej.nih.gov/ij/) software. The images were converted from red-green-blue (RGB) scale to gray scale (8 bit), threshold values adjusted to define the region of interest after subtracting the background value, and then calculated as integrated density. From the quantitative image analysis of fluorescence intensity, concentration-response curves were generated for E2, BPA and its analogues.

Statistical analyses were performed using GraphPad Prism 8.4.0. Dunnett’s test for significance of the observations was performed. ***P <.001, **P < .01, *P < .05 were considered statistically significant in our experiments.

### Treatments, RNA extraction, and quantitative real-time PCR

For qPCR, 20 embryos per well in a 6 well plate with 2 ml E3/well were exposed to vehicle only (DMSO 1% v:v), E2 (used as positive control) and a range of concentrations (10^-8^ M - 5x10^-6^ M) of BPA, BPAF, and BPC-Cl. The embryos were treated at day 4, and again at day 5 to boost up the expression. At the end of the exposure period at 6 dpf, the zebrafish embryos were homogenized in TRIzol reagent (Thermo Fisher Scientific). Total RNA from 20 pooled embryos were collected at 6 dpf using TRIzol and RNeasy spin columns (Qiagen) according to manufacturer’s protocol. DNAse I was used to remove any DNA in the column. RNA concentration was measured using BioTek Synergy H1 Hybrid Reader (BioTek Instruments, Inc.) and 1 μg of total RNA was used for cDNA synthesis using Superscript III reverse transcriptase with random hexamer primers (Applied Biosystems by Thermo Fisher Scientific).

The expression of *leptin, ghrelin, pck1, npy8ar, npy8br, trh, pmch, pmchlike, hcrt, bdnf, gcgb, pyy*, and *sim1a* was measured by quantitative PCR using a Real time PCR system (Applied Biosystems by Thermo Fisher Scientific) with PowerUp SYBER Green Master Mix (Applied Biosystems by Thermo Fisher Scientific) according to the manufacturer’s protocol. Primer sequences (Integrated DNA Technologies, Inc.) are shown in Table S1. Expression levels of *leptin, ghrelin*, and *pck1* were normalized to zebrafish *β-actin, EF1α* and *Rpl13α* by the ΔΔCt method (relative fold change expression of the genes of the sample). Expression levels of *npy8ar, npy8br, trh, pmch, pmchlike, hcrt, bdnf, gcgb, pyy*, and *sim1a* were normalized to zebrafish *EF1α* and *Rpl13α*). At least two biological experiments with 3 technical replicates each were used for calculation of statistical significance using Dunnett’s test for significance of the observations. *\*\*\*P < 0*.*001, **P < 0*.*01, *P < 0*.*05* were considered statistically significant.

## Results

### BPA and its analogues increase food intake in zebrafish larvae

We used zebrafish larvae in nacre background to assess their eating behavior after exposure to the bisphenols and E2. We treated the larvae at day 4 and day 5 with E2, BPA, BPE, BPAF, BPC, BPC-Cl, and BPS at increasing concentrations and fed them stained egg yolk powder for an hour on day 6. The larvae were imaged and the food in the gut was quantified by imageJ as integrated density (Figure 1A).

**Figure 1.**
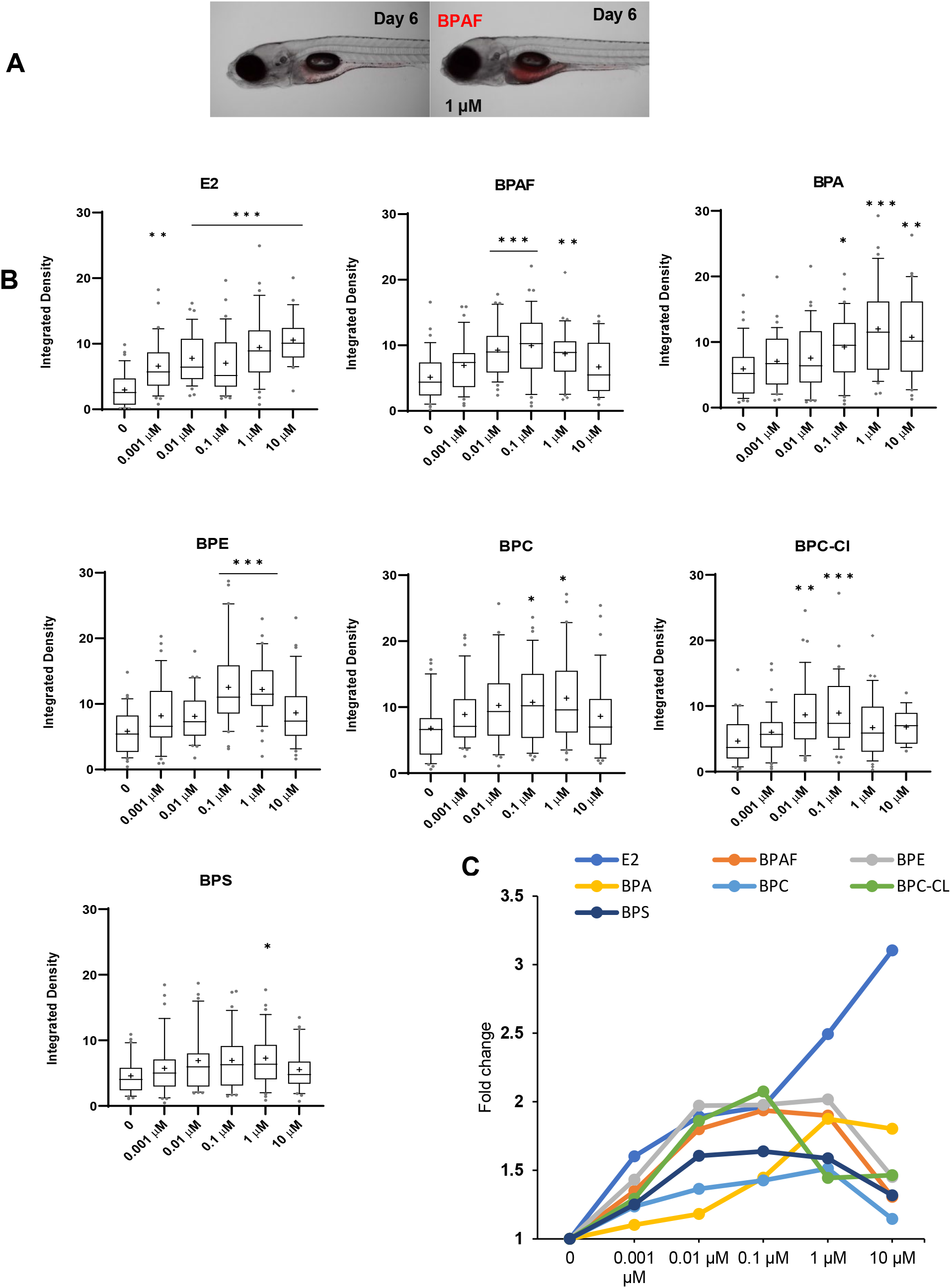
Bisphenol exposure increased feed consumption in zebrafish larvae. **A**. Left: Lateral view of control larvae. Right: Larvae treated with BPAF at 1 μM, showing food in the gut as fluorescence (red). Scale bar: 800 μm **B**. Quantification of food consumption a after bisphenol exposure. Larvae treated with E2 and bisphenols at 4 and 5 dpf for 48 h and fed for 1 h. Larvae exposed to increasing concentrations of E2, BPA, BPE, BPAF, BPC, BPC-Cl and BPS, as indicated. Integrated density values were measured using imageJ and analyzed by GraphPad Prism (GraphPad Software Inc). Each box represents 90% of the values; the ‘+’ sign indicates the average of the total value and the ‘-’ in the box indicates median or 50^th^ percentile of the values. Data were analyzed for significant differences using Dunnett’s test (treatment vs. control) (\*\*\**p* < 0.001, \*\**p* < 0.01, \**p* < 0.05). **C**. Comparison of potency of the different bisphenols and E2 using the average fold change expression as determined in B.

E2 treatment induced feed intake at the lowest concentration tested (0.001 μM) and more feed was eaten with increasing exposure concentrations (from 0.01 μM to 10 μM) (Figure 1B). Exposure to BPA and its analogues increased eating behavior in a concentration dependent manner. Quantification of feed amount showed a strong statistically significant (p<0.001) increase at exposures to 0.001 μM E2, 0.01 μM BPAF and BPC-Cl, and 0.1 μM BPA, BPE, and BPC compared to vehicle control. BPS exposure induced significant feed intake at 1 μM. At 10 μM concentration, the feeding started to decrease after exposure to all compounds except E2 and BPA likely because of toxicity. The feed intake after BPA exposure plateaued at 10 μM, whereas the feed intake after 3 μM E2 exposure further increased. Comparing the average fold change induction at 1 μM, we conclude that, E2 treatment induced the strongest feed intake followed by BPE, BPAF, BPA, BPS, BPC, and BPC-Cl (Figure 1C).

In each treatment group there was quite a bit of variation in the amount of feed in the gut in different larvae; some ate a lot and others very little. To analyze the collective eating behavior after exposure to the bisphenols, we determined the proportion of larvae that ate more than vehicle control larvae. After treatment with 0.001 μM E2, 80% of larvae had more feed in the gut than the average vehicle control larvae (Fig. 2). In larvae treated by the same concentration with BPAF, BPE, BPA, BPC-Cl, BPC, and BPS, 66%, 60%, 59%, 58%, 57%, and 48%, respectively, ate more than the average vehicle control. At 10 μM concentration, the highest proportion of larvae with large amounts of feed was observed for E2 and BPA where 96% and 74%, respectively, of larvae ate more than the control larvae. At this concentration, exposure to other bisphenols decreased eating. At 0.01 μM concentration, BPAF and BPC-Cl had the largest proportion (85% and 84% respectively) of larvae with more feed than the control. For BPE, BPC and BPS, the highest proportion (94%, 74%, and 70%, respectively) of larvae with more feed than the average controls were at 1 μM concentration.

**Figure 2.**
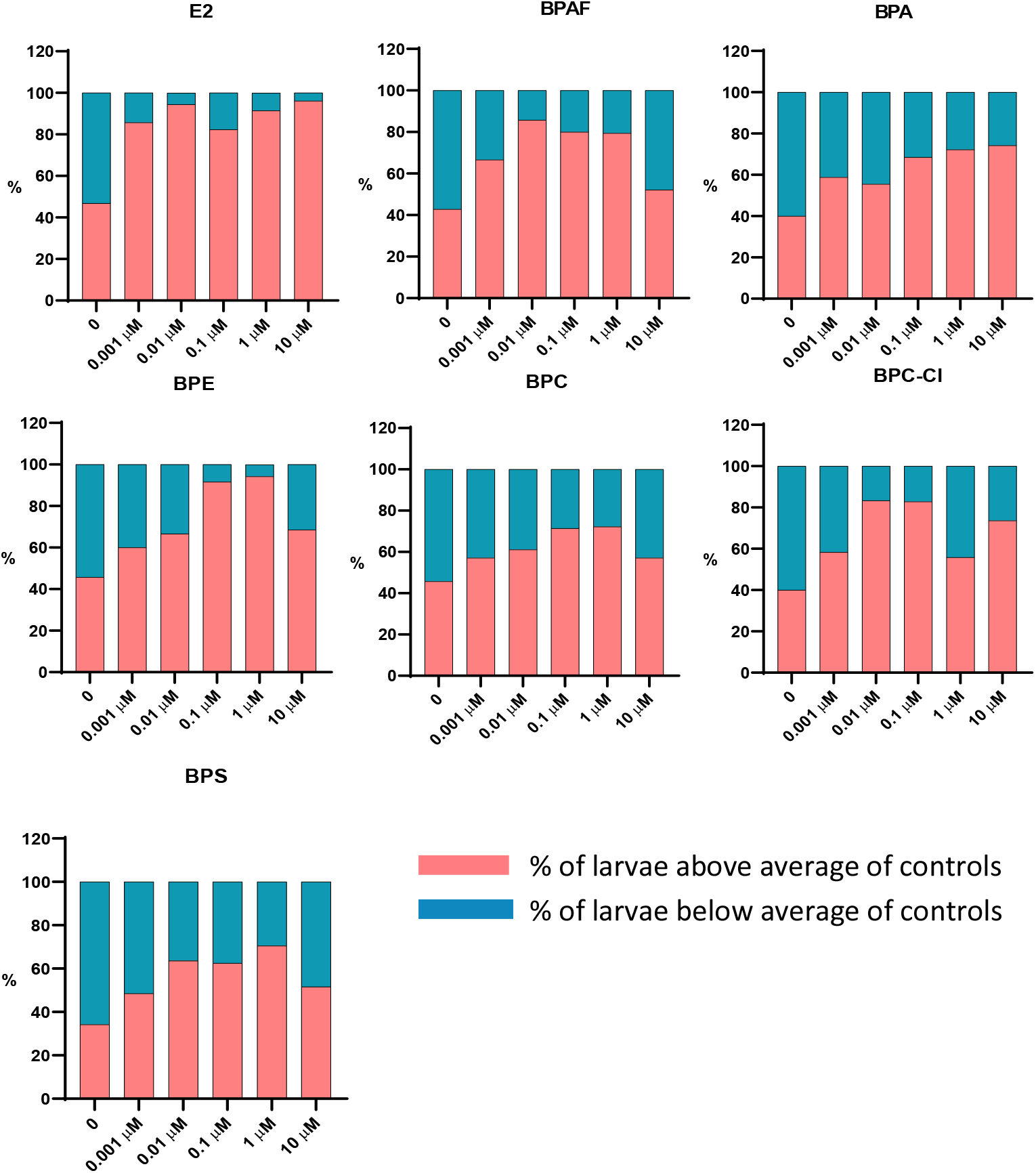
Proportion of larvae of the whole treated group having more or less feed in the gut than the untreated control larvae. Data from three biological experiments with 12 embryos/experiment were pooled together to calculate the percentage of larvae with more or less feed intake than the average feed intake in controls.

### Hunger and satiety gene activation by bisphenols in vivo

We selected BPAF and BPC-Cl to investigate the effects of exposure on the expression on appetite control genes by quantitative PCR because these compounds induced eating behavior at a low concentration (0.01 μM). The gene expression induced by BPAF and BPC-Cl exposure was compared to BPA and E2. The larvae were treated by repeated exposure at 4 and 5 dpf and RNA were prepared and analyzed at 6 dpf. The expression of the known hunger and satiety genes *leptin* and *ghrelin* was investigated together with the fasting-inducible gluconeogenic gene phosphenolpyruvate carboxykinase (*pck1*) (34).

Exposure to BPA, BPAF and BPC-Cl significantly decreased the expression of the satiety hormone *leptin* (Figure 3). *Leptin* expression was repressed by BPA and BPAF at 0.01 μM and by BPC-Cl at 0.1 μM. *Ghrelin* expression was significantly increased by BPC-Cl exposure at 0.1 μM. A similar trend was seen with BPAF exposure at 0.1-1 μM but it did not reach statistical significance. E2 treatment did not alter the expression of leptin and ghrelin.

**Figure 3.**
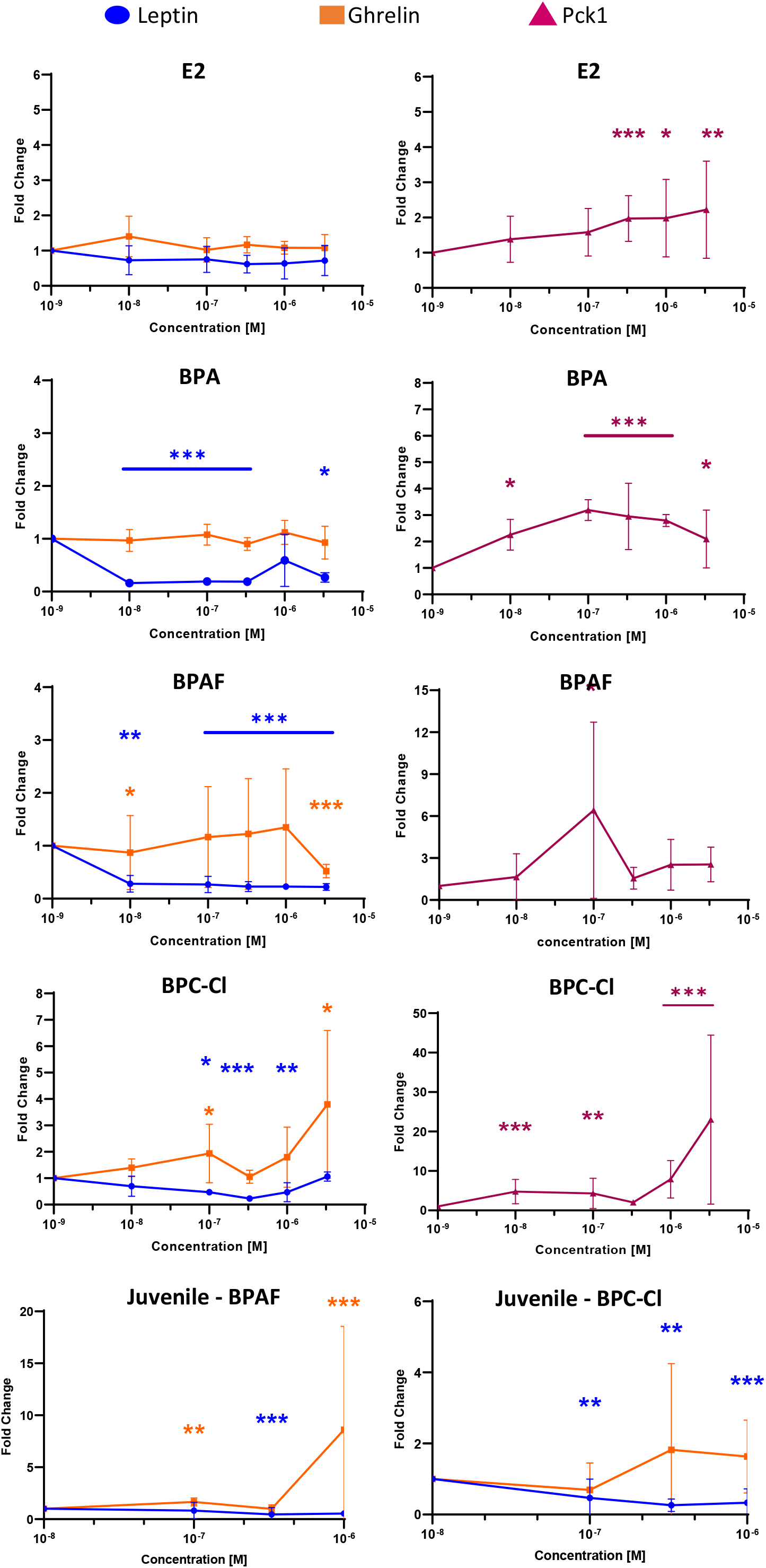
Expression of the appetite control genes *leptin* and *ghrelin*, and the gluconeogenesis marker *pck1* after exposure to E2, BPAF, BPA and BPC-Cl. qPCR after treatment of zebrafish larvae or juveniles with the bisphenols and E2. Transcript levels were normalized to an average of *β-actin, EF1α* and *Rpl13α* using ΔΔCT method. Data were analyzed for significant differences using Dunnett’s test (treatment vs. control) (\*\*\**p* < 0.001, \*\**p* < 0.01, \**p* < 0.05).

All the bisphenols and E2 activated expression of expression of *pck1* (Figure 3). *Pck1* expression was activated 2-fold at 0.3 μM E2. BPA exposure at 0.01 μM activated *pck1* expression. After BPC-Cl exposure, the expression of *pck1* was induced from about 5-fold at 0.01 μM to about 23-fold at 3 μM.

Because the zebrafish gut at 4 to 6 dpf is immature, we also analyzed the effect of BPAF and BPC-Cl exposure on the expression of appetite control genes in juvenile fish. These fish were treated by repeated exposure at 28 and 29 dpf and RNA was prepared and analyzed at 30 dpf. The expression of the known hunger and satiety genes *leptin* and *ghrelin*. BPAF significantly decreased *leptin* expression at 0.5 μM and significantly increased *ghrelin* expression at 9-fold at 1 μM (Figure 3). After BPC-Cl exposure, there was a tendency of increased *ghrelin* expression at 0.5 μM although not statistically significant. *Leptin* expression was significantly repressed by BPC-Cl from 0.01 μM to 1 μM.

We conclude that although there was some variation in the gene expression changes in larvae, bisphenol exposures generally resulted in an activation of *pck1* and *ghrelin* expression and a repression of *leptin* expression. The *in vivo* regulation of these appetite control genes was in accordance with the effects seen in eating behavior and indicated an increased hunger and decreased satiety in zebrafish larvae after exposure to the bisphenols.

We further explored 10 additional appetite-regulating genes in the zebrafish gut and brain because studies have shown that these genes have altered expression under various feeding circumstances (Table 1). Larvae were exposed to 1 μM of BPAF and BPC-Cl from 4 to 6 dpf and the gene expression was analyzed by qPCR. The larval samples were generated as previously described but only three samples were analyzed (control and 1 μM of each bisphenol).

**Table 1.** Appetite control genes in fish.

| Genes |  |  | Reference |
| --- | --- | --- | --- |
| Anorexigenic | *pmch* | Pro-melanin concentrating hormone | (35, 36) |
|  | *pmch-like* | Pro-melanin concentrating hormone -like | (35, 36) |
|  | *gcgb* | Encodes both glucagon and GLP-1 | (37) |
|  | *pyy* | Prepro-peptide yy | (38) |
|  | *sim1a* | Single-minded homolog 1-a | (39) |
| Orexigenic | *npy8ar* | NPY receptor a | (40) |
|  | *npy8br* | NPY receptor b | (40, 41) |
|  | *trh* | Thyrotropin releasing hormone | (42) |
|  | *hcrt* | Hypocretin neuropeptide precursor | (43, 44) |
|  | *bdnf* | Brain-derived neurotrophic factor | (39) |

Exposure to one or both of BPAF and BPC-Cl decreased the expression of the anorexigenic *pmch, pmchlike, pyy, sim1a* markers (Figure 4). *Gcgb* expression tended to be decreased but the repression was not statistically significant. On the contrary, the same compounds increased the expression of the orexigenic *npy8ar, npy8br*, and *trh* genes (Figure 4). The repression of *hcrt* and *bdnf* was not statistically significant.

**Figure 4.**
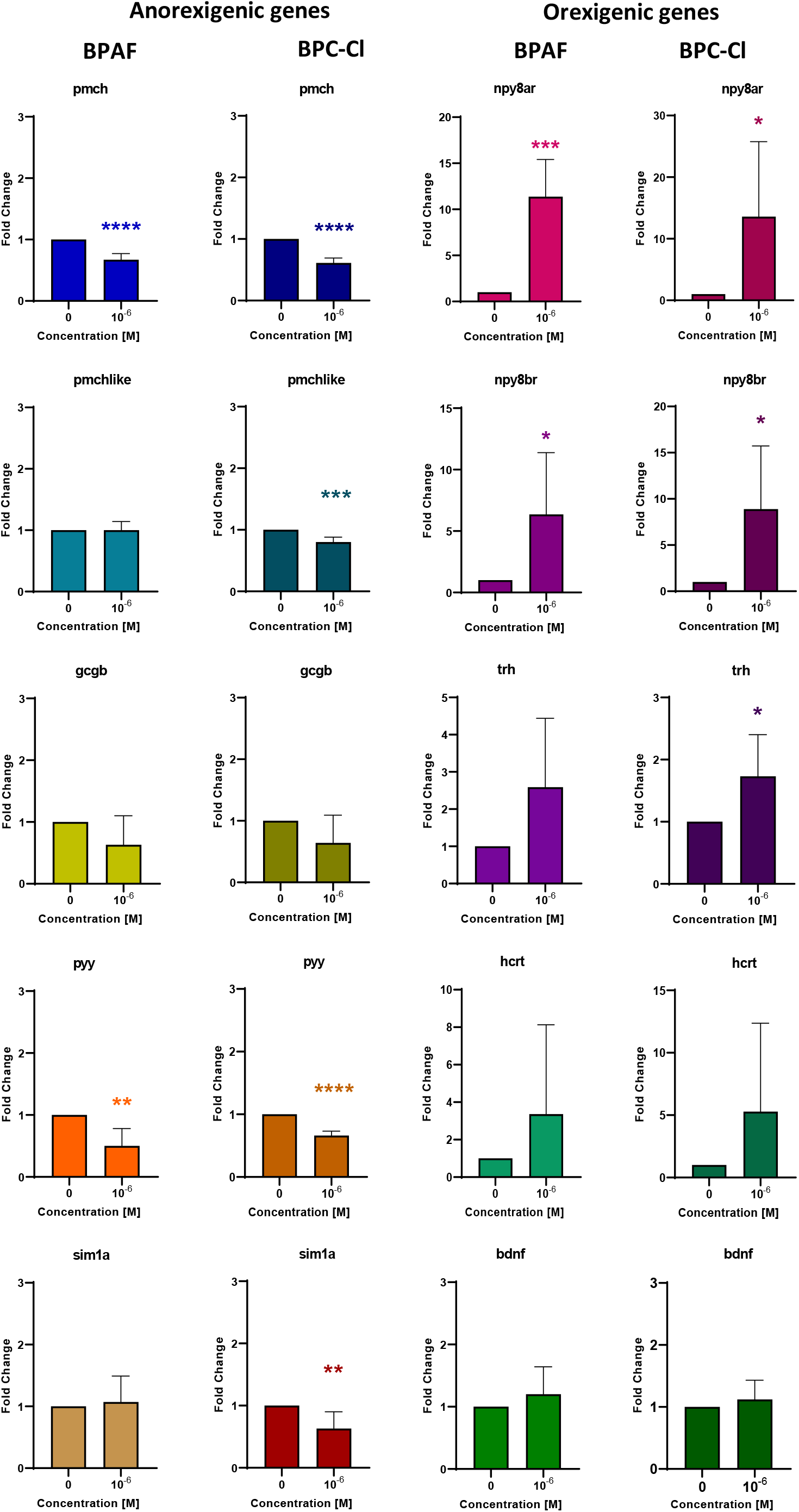
Expression of orexigenic and anorexigenic genes after exposure to BPAF, and BPC-Cl. qPCR after treatment of zebrafish larvae with compounds from 4 to 6 dpf. Transcript levels were normalized to an average of *EF1α* and *Rpl13α* transcripts using ΔΔCT method. Data were analyzed for significant differences using student’s t test (treatment vs. control) (***p < 0.001, **p < 0.01, *p < 0.05).

## Discussion

In this study, we demonstrated that BPA analogues are obesogenic chemicals in zebrafish. We developed an eating assay for 6 dpf zebrafish larvae consisting of ORO-stained egg yolk, available for consumption for one hour, followed by fluorescence imaging. Using this assay, we showed that BPE, BPC-Cl and BPAF were more potent in stimulating eating than BPA, whereas BPC and BPS were less potent than BPA. We further showed that although there is individual variation between larvae in how much feed they consume, bisphenol exposure increased the proportion of larvae with more food in their gut than the unexposed control larvae. For the eating assay, the bisphenols induced so called inverted “U-shaped” concentration response curves where feed consumption was first induced, but then declined at higher concentrations. This type of nonlinear relationship between a drug or chemical with an endpoint has been reported for many biological or pharmacological effects (45). The declined response in our study likely reflects the toxicity of the bisphenols making the larvae not eating at the higher exposure concentrations. For non-toxic E2, the eating did not decline, not even at the highest concentration tested. Thus, in addition to demonstrating that bisphenol exposures increased feed intake in zebrafish, our study indicates that zebrafish is as appropriate alternative *in vivo* model to score for eating behavior.

We further investigated the effect of bisphenol exposure on the expression of hunger and satiety genes. We found that the bisphenols in general activate transcription of the *ghrelin* and *pck1* genes. PCK1 mediates the first step of gluconeogenesis and it is a rate limiting enzyme in the process. If *pck1* expression is increased in zebrafish larvae, more glucose will be produced whether there is a need for it or not. This may in the end lead to an adverse effect on insulin production and metabolic disruption. At the time for our analysis (6 dpf), zebrafish larvae still absorb their lipids and nutrients from the yolk, and *pck1* expression is generally low (34, 46). Thus, this timepoint works well for investigating *pck1* inducing agents.

The increased *ghrelin* expression, in particular by BPC-Cl exposure, may indicate an induced appetite in the zebrafish larvae. In addition, *leptin* expression was decreased. Low levels of leptin hormone would cause the fish to eat more. Leptin is normally secreted by adipocytes, but at our early exposure time point (4-6 dpf), adipocytes have not yet been formed (47). This means that *leptin* likely was expressed in another tissue or cell type, such as adipocyte precursor cells. Nevertheless, the activation of *pck1* and *ghrelin* expression in combination with a decreased *leptin* expression together likely contributes to an adverse phenotype. To further investigate how bisphenols act on appetite control, we tested ten appetite regulating genes. We found that BPAF and BPC-Cl exposure increased the expression of several orexigenic markers (*npy8ar, npy8br, trh, hcrt*, and *bdnf*) and decreased the expression of anorexigenic markers (*pmch, pmchlike, gcgb, pyy*, and *sim1a*), thus further suggesting that exposure to bisphenols increase appetite in larval zebrafish.

We have previously reported on obesogenic effects of two bisphenol analogues, tetrachlorobisphenol A (TCBPA) and tetrabromobisphenol A (TBBPA), which induce lipid accumulation in larval zebrafish and late-onset weight gain in juveniles (32). Others have shown obesogenic effects of bisphenols. For instance, the size of adipocytes increases, and a larger body fat percentage is evident with age in rodents that were exposed to BPA during fetal development (21, 48). The model obesogen, tributyltin (TBT), also induces lipid accumulation in zebrafish larvae [38]. In mice, TBT exposure during fetal development increases the adult weight in the second and third generation of the progeny (49, 50). Increasing evidence suggests that BPA may have transgenerational effects by different mechanisms and interfere with female reproduction by influencing various epigenetic processes (51-54). For example, exposure to BPA could be transmitted transgenerationally by affecting female reproductive efficiency (55).

The potency of the bisphenol analogues in inducing feed consumption correlated with their estrogenic potency. We and others have shown that BPA and its analogues alter the estrogenic activity of estrogen reporter cells and estrogen reporter fish (25-27) and rodent models. For example, treatment of female rats with 50 and 100 mg/kg/day of BPA or its chlorinated derivatives cause a significant increase in the uterine wet weight and the endometrial area (56, 57). In our previous study, bisphenol analogues were ranked based on estrogenic activity scored in reporter fish from most to least estrogenic (E2, BPAF, BPE, BPC-Cl, BPC, BPA, and BPS) (25). For induction of feed consumption, the bisphenols grouped in a highly potent group (BPC-Cl, BPE, BPAF, and E2) and a less potent group (BPS, BPC, and BPA). Furthermore, similar concentration response curves were seen with BPC-Cl as it strongly activated both estrogen response and eating behavior at low concentrations, but at higher concentrations there was a rapid decline. This decline correlates to the antagonistic effect of BPC-Cl on the zebrafish estrogen receptors at higher concentrations (25). Also on human estrogen receptors, BPC-Cl operates as both as an agonist and antagonist (58-60). We conclude that there is a strong correlation between estrogenicity and eating behavior caused by bisphenol exposure. However, several bisphenol analogues also exhibit agonistic and/or antagonistic activity on the human estrogen related receptor (ERR), mineralocorticoid receptor (MR), progesterone receptor (PR), androgen receptor (AR), and thyroid hormone receptors (TRs) (61-63), and thus, obesogenic effects mediated via these receptors cannot be excluded.

Bisphenols and numerous other environmental chemicals interfere with the programming of endocrine signaling pathways that are created during embryonic development and have negative effects that might not become apparent until much later in life, such as obesity and diabetes (64). Future studies to both identify obesogenic chemicals as well as decipher their mechanisms of action are needed to protect the public from adverse effects of exposure to obesogenic chemicals.

## Acknowledgements

We thank Vladimir Korzh for the gift of the fish line GW57.

## Supplemental Information

Table S1. Primer sequences for PCR.

