## Supplemental Information for "Bisphenol A and its Analogues Alter Appetite Control in Zebrafish"

**Table S1. Primer sequences for PCR**

| Name of Primer | Sequence (5' to 3') |
|---|---|
| **Leptin F** | AGC TCT CCG CTC AAC CTG TA |
| **Leptin R** | CAG CGG GAA TCT CTG GAT AA |
| **Ghrelin F** | TGT GTC TCG AGT CTG TGA GC |
| **Ghrelin R** | TCT TCT GCC CAC TCT TGG TG |
| **Pck1 F** | GGC GTG TGA TGT ACG TGA T |
| **Pck1 R** | CCC TCC TCT TTA GCG ATG CG |
| **Pmch F** | CTT CCC TTC AGA AGA CAC CCC ATC A |
| **Pmch R** | TCC CTC CTT TAA TTC CTG TGT CCG C |
| **Pmch-like F** | AGT CCC TCA TCT CAA CGC AC |
| **Pmch-like R** | GCC GAT ACA CTC TTC CCA CC |
| **Gcgb F** | AGA AAA CCA GCG AGA CGA CA |
| **Gcgb R** | CCT CCG TGT CTT GTA AGG GG |

| | |
|---|---|
| **Pyy F** | TGG GGA TGA CAC AGA GCA CAA AC |
| **Pyy R** | GGG AGG CAC AGG TGG GTT TAG |
| **Sim1a F** | CTG TAC ATT TCA GAA ACG GCG |
| **Sim1aR** | AAG TGT GAA TGA TAG GGC TGG |
| **Npy8ar F** | CCA GGT CTC CAG GCG ATT G |
| **Npy8ar R** | ACT AAG CCC ACT GCG ATG AC |
| **Npy8br F** | CCT CTC ATG CTC CGA CAT CC |
| **Npy8br R** | CTG CTA CGG CCA GGT ATG AG |
| **Trh F** | CAG AAC AGC GAG AAC GAT CAG CC |
| **Trh R** | CGC TCC ATC TCC ACC TCC G |
| **Hcrt F** | CTA CGA GAT GCT GTG CCG AG |
| **Hcrt R** | CCA AGA GTG AGA ATC CCG AC |
| **Bdnf F** | AGC ATC TGT TGG AGT GTG TG |
| **Bdnf R** | CAG CTC TCA TGC AAC TGA AG |
| **Beta-actin F** | TTG CCC CGA GGC TCT CTT |
| **Beta-actin R** | GTT GAA GGT GGT CTC GTG GAT |
| **Ef1a F** | CTG GAG GCC AGC TCA AAC AT |
| **Ef1a R** | ATC AAG AAG AGT AGT ACC GCT AGC ATT AC |

| | |
|---|---|
| **Rpl13a F** | TCT GGA GGA CTG TAA GAG GTA TGC |
| **Rpl13a R** | AGA CGC ACA ATC TTG AGA GCA G |
